## Supplementary Figures S1-S4 and Text S1 for "Transcriptome dynamics of *Pseudomonas aeruginosa* during transition from replication-uncoupled to -coupled growth"

This file contains:

Supplementary Figures S1-S4

Supplementary Text S1

Supplementary Tables S1 and S2 are provided as external excel files

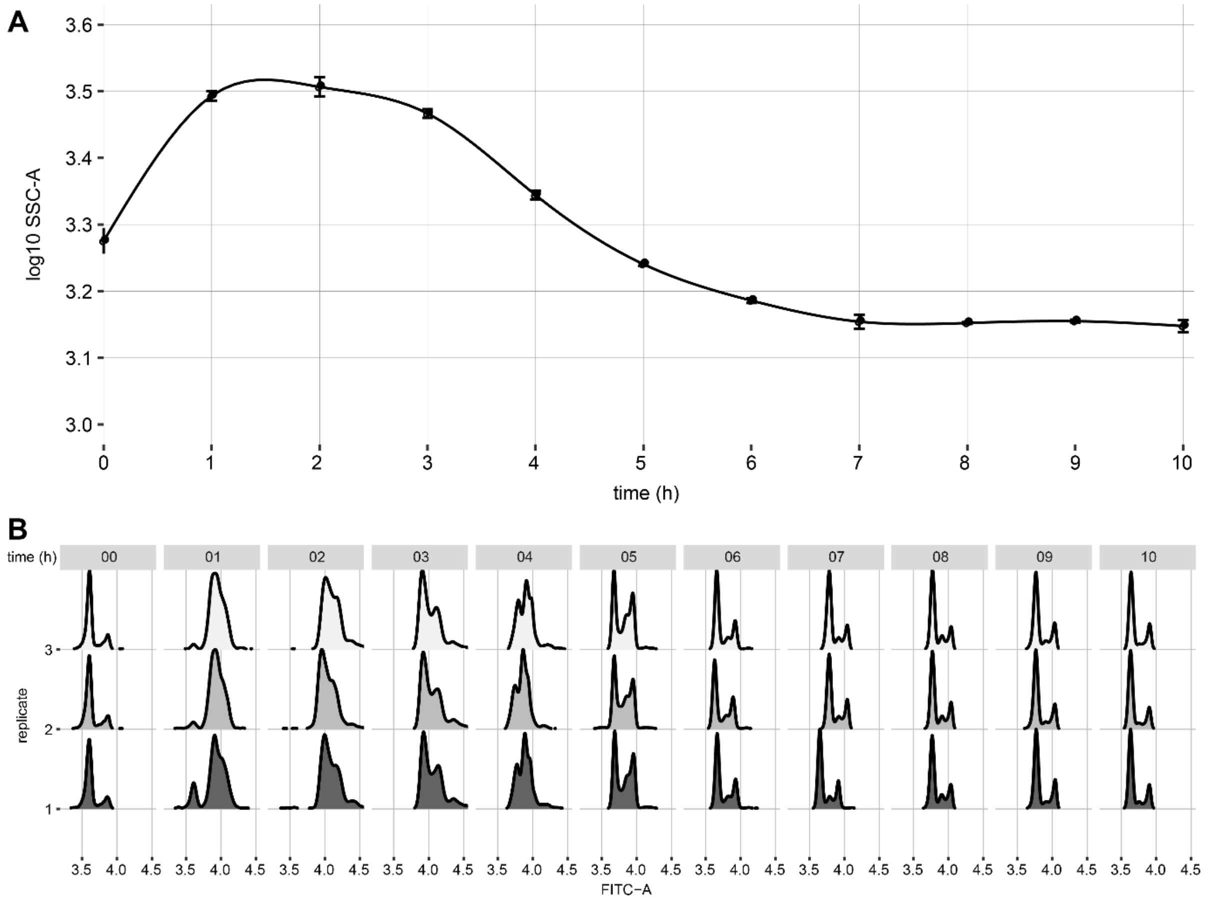


**Supplementary Figure S1. Flow cytometric determination of relative cell size and chromosome content during growth in LB medium. (A)** Changes of the side scatter (SSC) indicates reductive cell division from 3 h to 7 h cultivation time. **(B)** Changes in the distribution of chromosome content for three biological replicates in the course of 10 h cultivation.

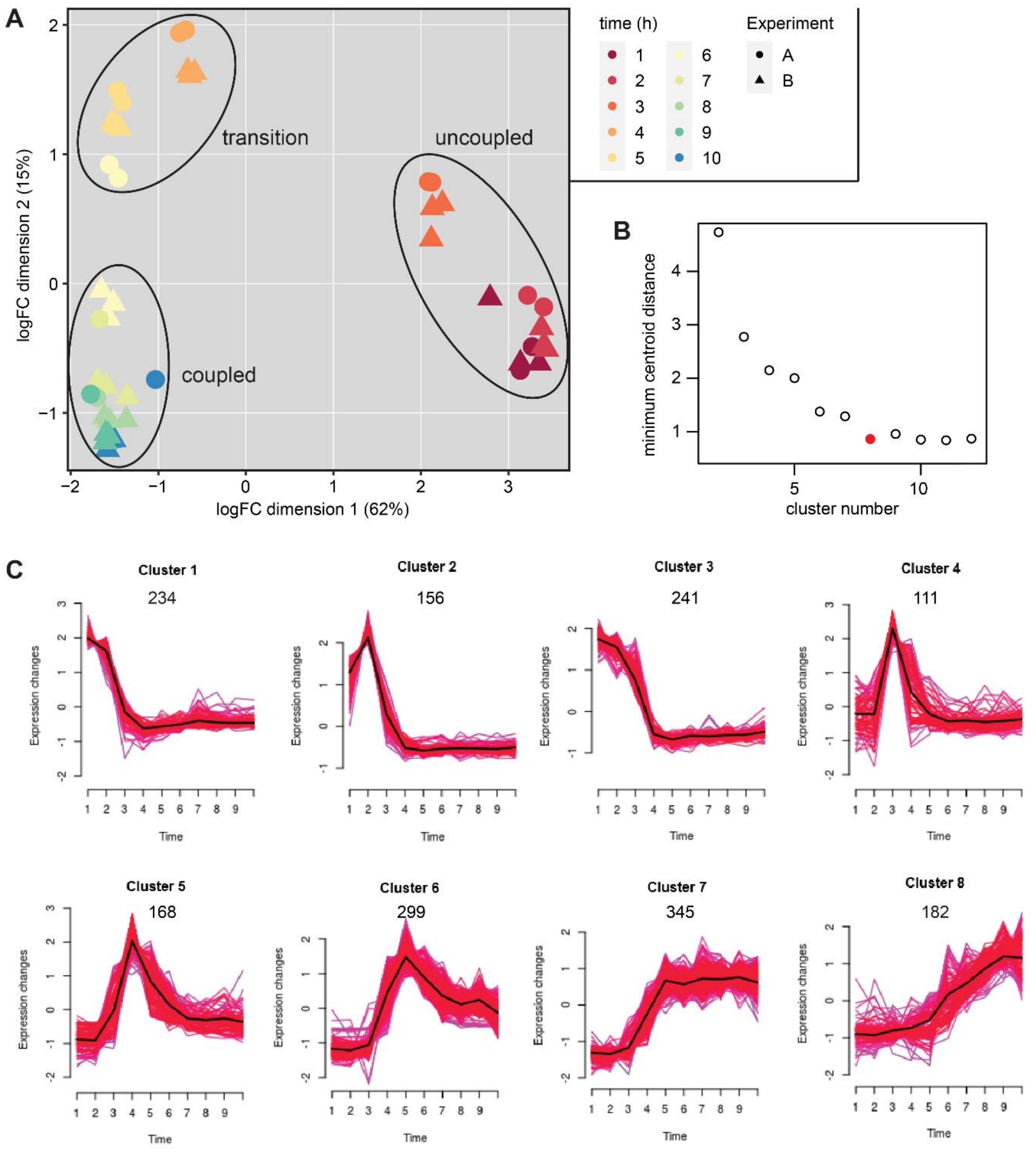


**Supplementary Figure S2. Transcriptome dynamics during growth in LB medium. (A)** Multidimensional scaling (MDS) plot of samples taken during 10 h cultivation. Note the different timing during the shift to coupled growth (6 h sample) for the two independent experiments. **(B)** Determination of ideal number of clusters based on the minimum centroid distance within the clusters. Increasing the number of clusters above 8 does not lead to further reduction of centroid distance. **(C)** Expression profiles of genes in the 8 clusters determined with the mfuzz-package. The number of genes within the cluster is shown below the cluster number. Cluster affiliation alongside expression data is also documented in Supplementary Table S1.

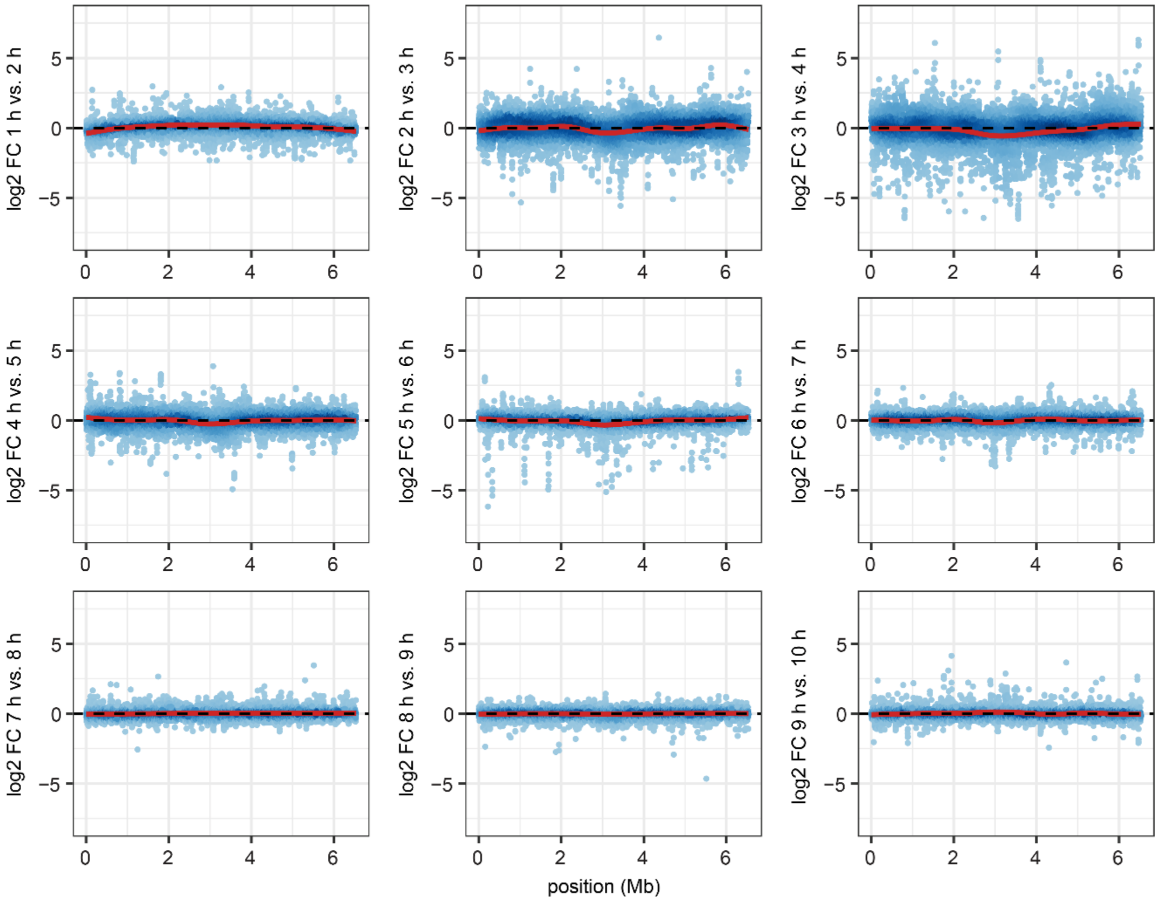


**Supplementary Figure S3. Time-resolved chromosomal gene expression changes during growth in LB medium.** $\log_2$ fold changes between subsequent time points are shown. Red lines show the fitted general additive models.

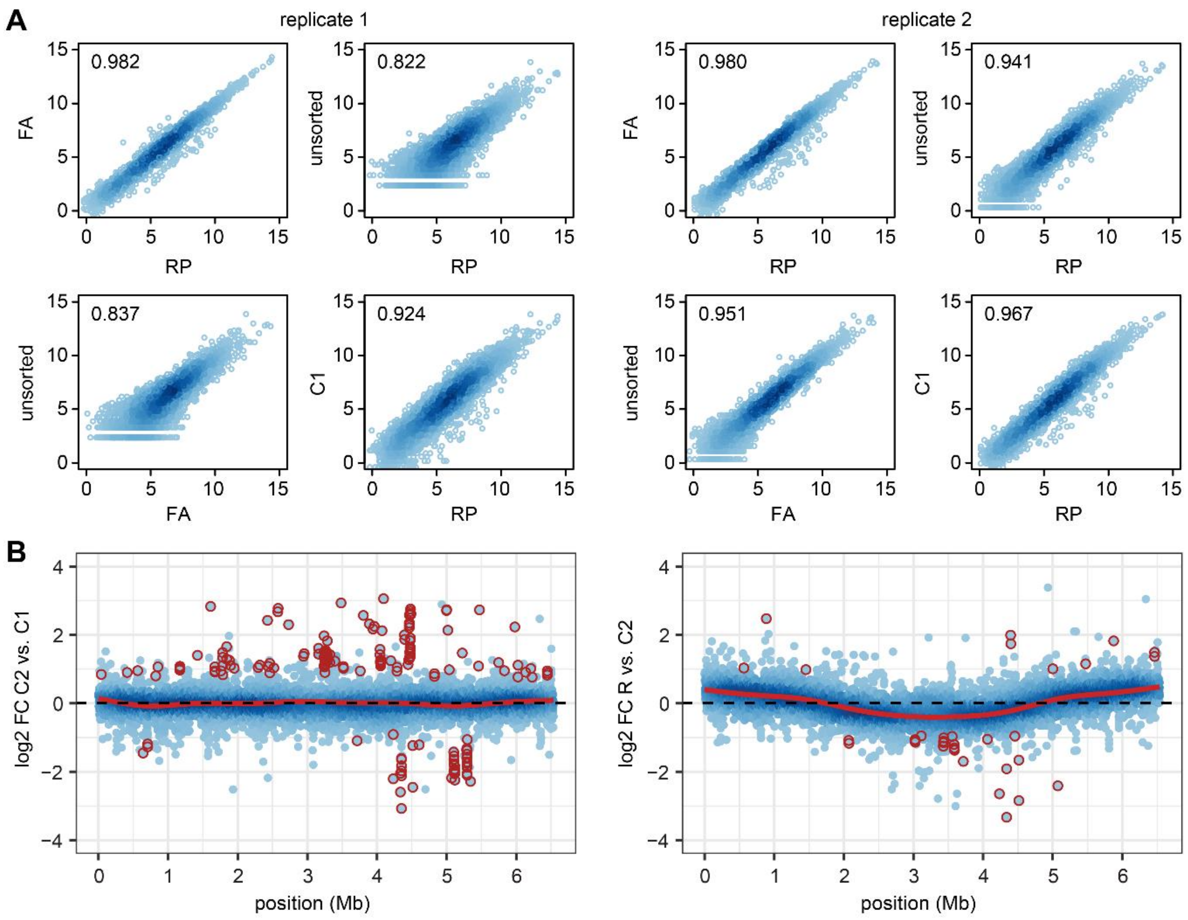


**Supplementary Figure S4. Transcriptomes of replicating and non-replicating cells during coupled growth. (A)** Correlation between transcriptomes of differently treated RNAs (see Figure 4A). **(B)** Chromosome-wide differential gene expression in replicating pre-divisonal (C2) versus non-replicating (C1) and replicating (R) versus non-replicating cells. Genes that change significantly in expression are marked in dark red. The red line shows a fitted general additive model.

### Supplementary Text S1. FACS and RNA isolation for replication transcriptomics

**Samples to take**

- Fraction 1 ($C_{1n}$)
- Fraction 2 (replicating cells)
- Fraction 3 ($C_{2n}$)
- Fixed, unsorted, went through FACS
- Fixed, but unsorted
- Unfixed, RNA protect, unsorted

**Day 1 - Plating of strain**

1. Streak an LB plate with PA14 wt from glycerol stock
2. Keep at 37°C overnight

**Day 2 - Overnight culture (early afternoon)**

1. Collect many colonies by streaking the plate with an inoculation loop
2. Resuspend in 1 mL LB in a 1.5 mL tube
3. Inoculate a 50 mL flask with 10 mL LB with 80 µL bacterial suspension
4. Keep in an incubator at 37°C shaking at 180 rpm

**Day 3 - Culture growth**

1. Inoculate 50 mL LB in a 250 mL flask at $OD_{600}$ 0.2
2. Cool centrifuge to 4°C
3. Put 4% formaldehyde in PBS on ice
4. After 5 h measure $OD_{600}$, early stationary phase should be reached ($OD_{600}$ 2 equals ca. $1.8*10^9$ cells/mL)
5. RNA Protect sample
    - Mix 1 mL culture with 1 mL RNA Protect bacterial reagent (QIAGEN) in a 2 mL tube
    - Leave for 10 min at RT
    - Pellet at 8000 rpm for 5 min
    - Freeze at -70°C for later RNA isolation
6. Fixation with formaldehyde (duplicates)
    - Pipette 1 mL culture in a 1.5 mL tube
    - Pellet cells at 8000 rpm for 2 min at 4°C
    - Resuspend in 1 mL ice-cold 4% formaldehyde in PBS
    - Shake in a rotator at 4°C for 2 h
    - Pellet cells at 8000 rpm for 2 min at 4°C
    - Resuspend in 1 mL PBS + 0.01 U/µL SUPERase In RNAse Inhibitor (Thermo Fisher, stock 20 U/µL)
    - Pellet cells at 8000 rpm for 2 min at 4°C
    - Resuspend one pellet in 1 mL PBS + 0.01 U/µL SUPERase In RNAse Inhibitor
    - Put in fridge for sorting on next day
    - Freeze pellet in other tube at -70°C for later RNA isolation

**Day 4 - Flow cytometry sorting**

**Staining**

1. Start with the staining ca. 45 min before the sorting
2. Dilute the culture 1:100 in a 50 mL falcon (300 µL culture + 27.7 mL filtered PBS = $1.8*10^7$ cells/mL, total $5.4*10^8$ cells)
3. Add 1 mL SYBR Green (Sigma, S9430, diluted to 100x)
4. Incubate for 20 min at RT, keep protected from light

**Sorting (ca. 9 h) on BD FACSAria Fusion**

Bring one bottle RNA Protect bacterial reagent and a bit SYBR Green for the control staining

1. Sort into RNA protect bacterial reagent
    - $3*10^7$ cells for each of the 3 fractions
    - $3*10^7$ unsorted
    - Sort 1000 cells of each fraction **not into RNA Protect** and restain with SYBR Green to control correct sorting

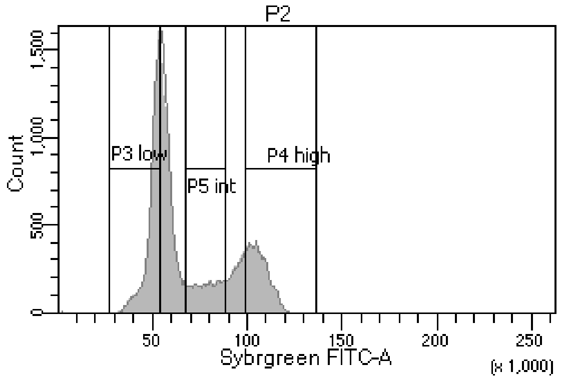


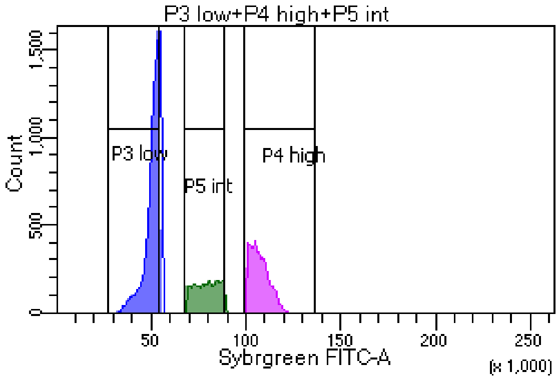


**Filtering**

1. Wash all devices used for filtering and isolating RNA with RNAseZap and EtOH
2. Prepare filtering device
3. Prewash each filter (0.22 µm, Millipore) with a few mL RNase-free water
4. Filter cells through it
5. Wash filter again with a few mL RNase-free water
6. Extract filter from the filtering device with tweezers
7. Cut with a clean, sterile scalpel, on a sterile petri dish, into pieces
8. Transfer into a 2 mL screw-cap tube
9. Freeze at -70°C until use

**Day 5 - RNA isolation of sorted samples**

Modified protocol from *https://www.nature.com/articles/s41598-018-32997-9*

1. Pre-treatment of filter for more efficient lysis with lysozyme:
    - Cover filter in 200 µL of 800 µg/mL lysozyme (8 µL of a 20 mg/mL Stock + 192 µL TE pH 8.0)
    - Vortex for 30 s
    - Incubate for 10 min at RT, vortex 2-3 times while incubating

2. Proteinase K digestion to reverse crosslinking from fixation (with QIAGEN DNeasy blood & tissue kit)
    - Add 180 µL ATL + 20 µL Proteinase K
    - Incubate at 56°C for 3 h
3. Add 500 µL NucleoZOL (Takara Bio) and mix
4. Add 100 mg beads (Zirconia beads, 0.1 mm + 2 mm diameter)
5. Shake in Fast-Prep Instrument (MP Biomedicals) for 3 x 1 min at 4.0 m/s and 30s at 5.0 m/s, put sample on ice in between each step for a minute
6. Add 200 µL RNase-free water
7. Shake in Fast-Prep for 15 s at 5.0 m/s
8. Incubate at RT for 15 minutes
9. Centrifuge for 15 min at 12,000 x g at RT
   A semi-solid pellet containing DNA, proteins and polysaccharides forms at the bottom of the tube. The RNA is still solubilized in the supernatant.
10. Transfer up to 700 µL supernatant to a fresh tube. Leave a layer of the supernatant above the DNA/protein pellet.
    The pellet containing DNA, protein, and polysaccharides comprises approximately 10% in volume of the total homogenate-water mix.
11. Add 3.5 µL (0.5% of supernatant volume) 100% 4-bromoanisole to the supernatant
12. Vortex for 15 s and incubate at RT for 5 min
13. Centrifuge for 10 min at 12,000 x g at RT.
    Residual DNA, proteins, and polysaccharides accumulate in the organic phase at the bottom of the tube. RNA is still solubilized in the supernatant.
14. Pipette RNA containing supernatant into a fresh tube
15. Add 700 µL of isopropanol per 700 µL supernatant in order to precipitate RNA
16. Add 2 µL Glycoblue (Invitrogen)
17. Incubate samples at -20°C overnight (-70 ° C for 1 h is also possible)
18. Centrifuge samples for 10 min at 12,000 x g at 4 °C.
19. Wash pellet twice with 500 µL ice-cold 70% ethanol and centrifuge for 5 minutes at 4 °C
20. Carefully let it dry a bit, but not too much, as this may lead to a decrease in solubility (2 - 5 min)
21. Resuspend pellet in 18 µL RNase free water
    To avoid having to use speed vac for decreasing the volume prior to rRNA depletion, resuspension of the low concentrated samples can be done in 14 µL (3 µL for Qubit, 11 µL for rRNA depletion)
22. DNase treatment (DNA-free Kit, Thermo), add
    - 0.1 volume 10X DNase Buffer
    - 1 µL rDNase I
    - Mix gently, incubate at 37 °C for 20-30 min
    - Add at least 2 µL (~0.1 volume) of Inactivation reagent
    - 2 min RT, mixing occasionally
    - 10,000 x g for 2 min
    - Transfer RNA containing supernatant into a fresh tube
23. Measure concentration on Qubit
    - Expected RNA yield for $3*10^7$ cells is 70 - 100 ng
    - In 18 µL → 4 - 5.5 ng/µL
    - Qubit detection limit: 5 ng in diluted sample
    - 3 µL (ca. 12 ng) for Qubit measurement

**RNA isolation of RNA Protect and FA fixed sample**

Modified QIAGEN QIAshredder & RNeasy Plus Kit protocol

1. Thaw pellet, centrifuge 5 min, 8000 rpm, remove supernatant
2. + 100 µL TE with lysozyme (4 µL of a 20 mg/mL stock + 96 µL TE), resuspend vortex 30 s; incubate 10 min RT, vortex 2-3 times, scratch off the cells from Eppi walls with a pipette tip
3. + 350 µL of **RLT** Buffer with Mercaptoethanol (10 µl Mercaptoethanol + 1mL RLT Buffer).
4. 60 min at -70°C.
5. Thaw, load into QIAshredder column, centrifuge 2 min, 14,000 rpm → save flow through!
6. Flow through +450 µL 70% EtOH, mix well
7. Load into RNeasy column (700 µL max.), centrifuge each time 20 s, 10,000 rpm; discard flow-through
8. + 700 µL **RW1**, centrifuge 20 s, 10,000 rpm
9. column into new 2 ml tube, +500 µL **RPE**, centrifuge 20 s, 10,000 rpm; discard flow-through
10. + 500 µL **RPE**, centrifuge 2 min, 14,000 rpm, discard flow-through
11. Centrifuge 2 min, 14,000 rpm, discard flow-through → save column (to new 1.5 tube)
12. Air dry 5 min, RT
13. + 50 µL $H_2O$, incubate 1 min, centrifuge 1 min, 10,000 rpm
14. DNase treatment (DNA-free Kit, Thermo Fisher):
    - 50 µL RNA
    - 5 µL DNase Buffer
    - 1 µL DNase
    - Mix gently and incubate for 20 min at 37°C, then add one more µL and incubate for additional 10 min at 37°C
    - +5 µL Inactivation reagent, vortex, incubation 2 min RT
    - Centrifuge for 2 min, 10,000 rpm
15. Transfer supernatant to new tube, measure the RNA concentration with Qubit
16. Store sample at -70°C

**rRNA depletion with NEBNext® rRNA Depletion Kit (Bacteria), #E7850L**

1. Dilute the high concentrated samples (RNA Protect samples) to 1 µg/11 µL in RNase-free water
2. Use Speed Vac to reduce the volume of the low concentrated sample to 11 µL
3. Follow protocol as described in manual
4. Resuspend RNA in 12 µL water, high concentrated in 17 µL water
5. Use 1 µL of high concentrated samples for Nanodrop measurement (ca. 5% of RNA should be left)

**Library preparation and RNAseq**
